# Alternative splicing reshapes protein interaction networks through structured, directional rewiring across cancers

**DOI:** 10.64898/2026.06.24.734271

**Authors:** Jeffrey Zhong, Ruth Dannenfelser, Vicky Yao

## Abstract

Alternative splicing generates extensive transcript diversity in cancer, but whether the resulting changes to protein-protein interaction (PPI) networks are structured across tumors or idiosyncratic to individual patients remains unresolved. Existing analyses catalog splicing events and their effects on individual interactions, but do not reconstruct patient-specific interaction networks at scale, leaving the systems-level architecture of splicing-driven rewiring uncharacterized. Here, focusing on exon skipping, the most prevalent form of alternative splicing in humans, we analyze 7,950 tumors across 28 cancer types to reconstruct patient-specific rewired PPI networks relative to matched normal tissues, distinguishing interaction gains from losses. We identify widespread and structured remodeling of protein connectivity, including a unique pattern of interaction gains in brain cancers, a conserved pan-cancer axis of interaction loss that converges on GTPase and Ras signaling, and a subset of genes that function as bidirectional switches, where rewiring direction varies across cancer types. At the patient level, tumors exhibiting extreme rewiring, especially pronounced interaction gains, are associated with significantly poorer survival, independent of tumor stage and mutational burden. These results delineate systems-level patterns through which alternative splicing reshapes protein interaction networks and contributes to tumor heterogeneity across cancers.

## Introduction

Alternative splicing is a pervasive source of transcript diversity in human tumors, with cancer cells exhibiting up to 30% more alternative splicing events than their normal tissue counterparts [1]. Large-scale pan-cancer analyses have made substantial progress in characterizing these alterations, cataloging thousands of tumor-specific splicing events and neojunctions [1], mapping their effects on protein domain architecture and interaction interfaces [2, 3], and identifying associations between somatic variants and altered splicing programs [1, 3]. In parallel, studies of individual genes have demonstrated that specific aberrant transcripts can promote proliferation [4], suppress apoptosis and DNA repair [5], enable immune evasion [6], drive angiogenesis [7], and confer drug resistance [8].

Yet even the most comprehensive of these analyses assess the functional impact of each splicing alteration independently, without modeling how thousands of concurrent changes collectively rewire protein interaction network topology within individual tumors. Such rewiring could, in principle, be structured and convergent across tumors, or it could be heterogeneous and patient-specific. Resolving this is important because the number of splicing events in a tumor is inversely correlated with the burden of somatic driver mutations [3], raising the possibility that splicing-driven network remodeling represents an axis of tumorigenesis that may at times converge with somatic mutation and at others act independently of it. The downstream impact and clinical significance of systems-level changes due to splicing remain to be thoroughly explored.

Protein-protein interactions (PPIs) offer a natural framework for addressing this question, because alternative splicing directly modulates the domain composition of proteins and can thereby disrupt existing interactions or create new ones [9]. Several computational approaches have been developed to model these effects, from domain-based prediction of isoform interactomes [10] to interaction maps constructed at the transcript [11, 12] or exon level [13–15]. Compared to other exon-level methods, our previously developed tool Splitpea [15] is able to calculate both differential exon usage relative to a background sample set and infer the directionality of PPI changes, separating interaction gains from losses, giving a more complete snapshot of protein rewiring. A recent meta-analysis of splicing induced protein changes examined individual PPIs affected by exon-altering events across 15 tumor types and 684 samples [16], but did not reconstruct networks at the patient level, leaving the systems-level architecture unaddressed. More broadly, while existing tools and studies have mapped splicing changes to individual protein interactions, none have systematically reconstructed patient-specific rewired PPI networks at scale to determine whether splicing-driven interaction changes exhibit structured, network-level organization within and across cancer types.

Here, we present a systematic analysis of splicing-driven PPI network rewiring across 7,950 cancer samples spanning 28 cancer types from The Cancer Genome Atlas (TCGA) [17], using matched normal samples from 20 different tissues in the Genotype-Tissue Expression (GTEx) project [18] as reference (Figure 1). We focused on exon skipping events, the most prevalent form of alternative splicing in humans [19], to capture a large and mechanistically interpretable segment of splicing variation. By applying Splitpea to each tumor sample, we reconstructed patient-specific rewired PPI networks that quantify the gain, loss, or mixed disruption of protein interactions relative to normal tissue. Our approach shifts from the standard view of cataloging individual splicing events to characterizing their aggregate functional impact on protein connectivity at the level of individual patients, cancer types, and the pan-cancer landscape.

**Figure 1.**
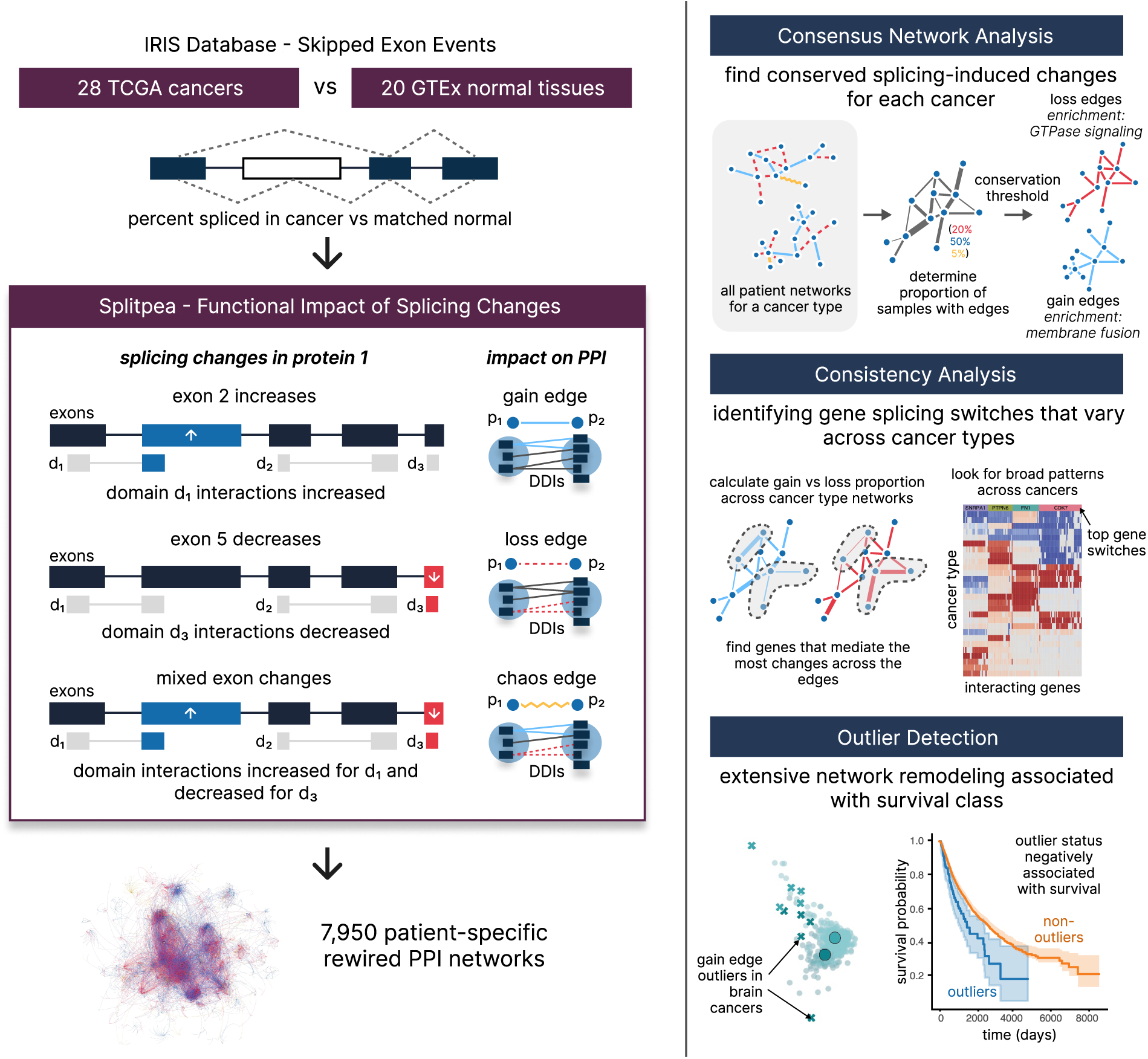
Using Splitpea to uncover unique aspects of splicing-based PPI rewiring within and across cancer types. (Left) For each individual cancer sample, a set of tissue-matched normal reference samples was used to filter SE events, retaining only those that were significantly different. These SE events were then integrated with a reference human PPI network using Splitpea, which maps SE changes onto proteins and their associated domain-domain interactions to predict alterations in network edges. This process was repeated for each patient in IRIS, yielding 7,950 patient-specific rewired PPI networks. (Right) Our rewiring analysis allows us to discover three unique insights: conserved splicing changes that represent core cancer and cancer-wide processes, gene switches responsible for large PPI network changes, and patient splicing outliers associated with prognosis.

Across cancer types, we find a conserved pan-cancer program of protein interaction loss that converges on the GTPase and Ras signaling pathways, with brain cancers representing a striking exception dominated by new protein interactions. We identify a set of highly connected genes, including both established cancer drivers and core splicing regulators, that function as context-dependent switches whose rewiring direction varies across cancer types. At the patient level, we find that tumors whose networks deviate most from the norm for their cancer type, particularly those with pronounced interaction gains, exhibit significantly worse survival outcomes independent of tumor stage and mutational burden. Drawing on our findings, we postulate that the network-level consequences of alternative splicing are an underexplored dimension of cancer biology, accounting for mechanistic effects beyond those established by mutational drivers.

## Results

### Tissue-corrected exon skipping landscapes across 28 tumor types

To characterize the landscape of splicing changes across cancer types, we analyzed tumor and normal tissue splicing data from the IRIS database [20]. This resource quantifies alternative splicing for 7,950 samples across 28 tumor types from The Cancer Genome Atlas (TCGA) [17] and 5,968 normal tissue samples from The Genotype-Tissue Expression Project (GTEx) [18]. On average, each cancer type includes approximately 284 patients, ranging from 36 (cholangiocarcinoma, CHOL) to 1,088 (breast cancer, BRCA). We focused on the percent spliced in (PSI or *ψ*) values for skipped exon events (SE), the most prevalent type of splicing event. Individual samples within a cancer type varied widely in total SE event counts (Figure 2A), with a median of approximately 25,000 events across all cancer types and several showing substantially more, including glioblastoma multiforme (GBM; median = 35,279.5), stomach adenocarcinoma (STAD; median = 36,876.5), ovarian carcinomas (OV; median = 39,205), and esophageal carcinomas (ESCA; median = 44,494). While these findings are consistent with previous reports [21], we noted that many of these events also occur in normal tissues and do not necessarily represent tumor-specific changes.

**Figure 2.**
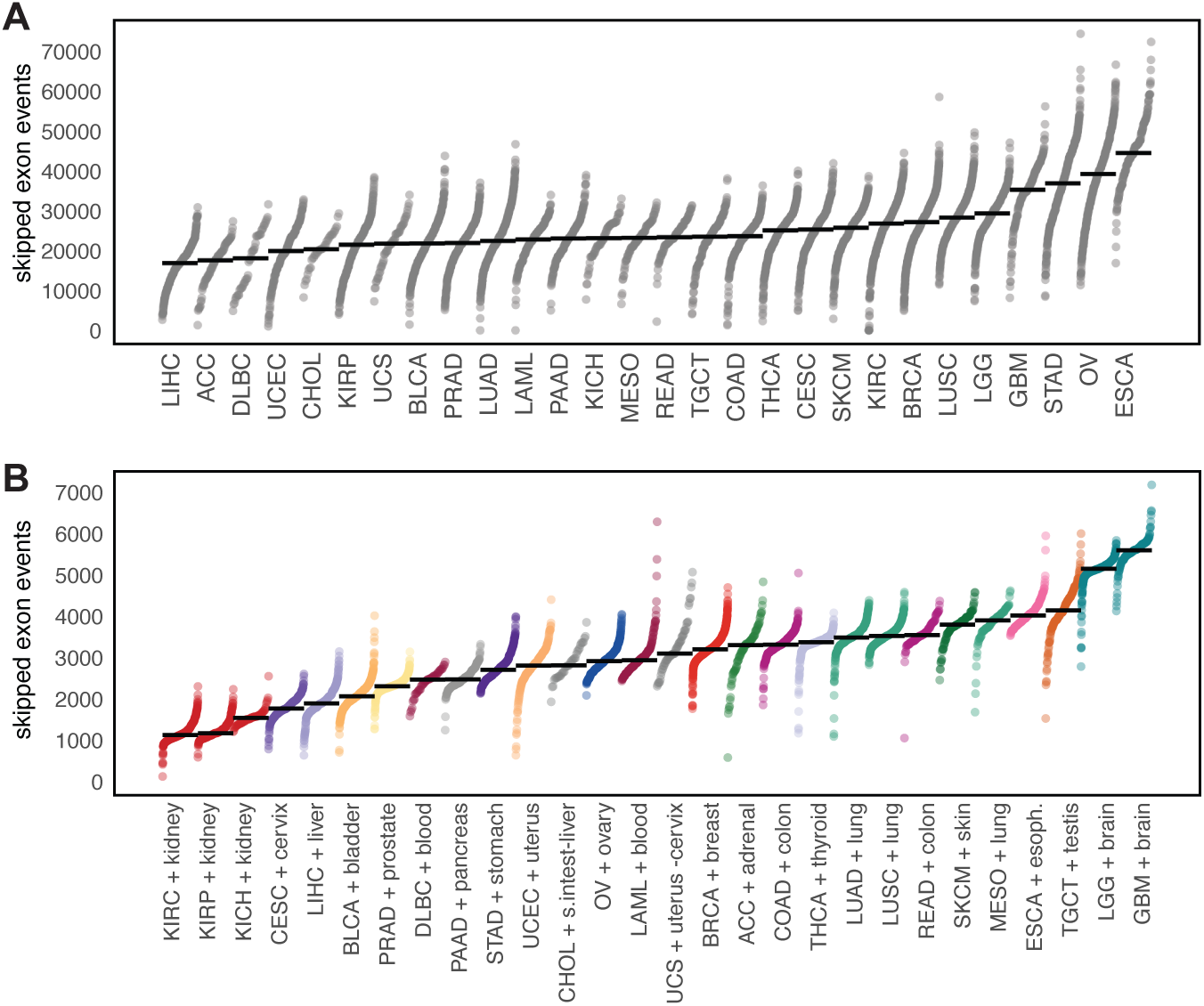
Skipped exon splicing event distributions across cancers. (A) Scatterplot of number of skipped exon events per sample for each cancer type before normal tissue correction. The y-axis shows the event count and the x-axis lists cancers ordered by their median event count; individual dots represent samples (ordered within each cancer by event count) and the line indicates the median value. (B) Scatterplot of the same metric after matching cancers to their corresponding normal tissue background and filtering to only significant splicing events. The colors represent matched normal tissues.

To account for background tissue variation, we matched each TCGA cancer type to its closest GTEx normal tissue (e.g., lung squamous cell carcinoma to lung tissue; Table S1) and retained only SE events with significant changes in PSI relative to the matched normal background (see Methods). For two cancer types, CHOL and uterine carcinosarcoma (UCS), we identified two plausible tissue matches each and used both as background. This filtering reduced the total number of SE events approximately tenfold and substantially altered the ranking of cancer types by median event count (Figure 2B). GBM, lower grade glioma (LGG), and testicular germ cell cancer (TGCT) emerged as those with the highest frequency of tumor-specific SE events, while the three renal cancers (KIRC, KIRP, KICH) had the fewest. This pattern is consistent with the known splicing complexity of their tissues of origin, especially in the brain and testis [22]. Furthermore, the shift in rankings after tissue correction, most pronounced in renal clear cell carcinoma (KIRC), which drops from 8th to last, highlights the importance of accounting for baseline tissue splicing when interpreting tumor splicing landscapes.

Interestingly, while most cancers sharing the same tissue background showed similar median event counts after correction, there were some notable exceptions. For example, colon adenocarcinoma (COAD) and rectal cancer (READ) diverged despite a shared colon background, consistent with well-documented clinical and molecular differences between these cancers [23]. Hepatocellular carcinoma (LIHC) and CHOL, as well as cancers of the lung and surrounding pleura, mesothelioma (MESO) and LUSC, also showed discordant patterns that may reflect suboptimal tissue matching or genuinely distinct pathogenic characteristics.

### Construction of 7,950 patient-specific rewired PPI networks reveals a pan-cancer asymmetry favoring interaction loss

To determine whether tumor-specific exon skipping events produce systematic changes in protein connectivity, we applied Splitpea [15] to all cancer samples, generating a rewired PPI network for each patient (Figure 1). Starting from a consensus human PPI network of known interactions, Splitpea integrates splicing quantification data with protein domain annotations to predict edge-level alterations. By aligning exon coordinates with protein domain annotations, it maps splicing alterations onto protein domains, thereby identifying disruptions in domain-mediated interactions. For each protein pair, the method maps SE changes onto the constituent protein domains and assesses their impact on the domain-domain interactions that mediate the PPI. This yields three categories of rewired edges: gain edges, where splicing changes increase domain-mediated interactions; loss edges, where they decrease domain-mediated interactions; and chaos edges, where interacting domains provide a mixed signal of both possible gain and loss, resulting in an ambiguous readout. Using matched GTEx normal tissue samples as reference (Table S1), we generated 7,950 rewired PPI networks spanning 28 cancer types. The direction of predicted splicing-driven rewiring tended to be concordant across both proteins in an interacting pair, with chaos edges being rare in most cancer types (Figure S1).

In 26 out of the 28 cancer types we analyzed, interaction losses substantially outnumbered gains (Figure 3A). Aggregating across all patient samples, 76.7% of rewired edges corresponded to potential interaction losses and 20.8% to potential gains. The magnitude of this asymmetry varied across cancers, with chromophobe renal cell carcinoma (KICH) showing the most pronounced loss bias (92.6% loss, 6.4% gain), while the two brain cancers were the only exceptions to the overall pattern. Lower grade glioma (LGG) and glioblastoma (GBM) showed much larger gain fractions relative to other cancers (56.5% and 48.8% of edges, respectively). We further examined the relationship between the number of tissue-corrected SE events per patient and the number of rewired edges (Figure 3B), since samples with high proportions of gain or loss edges relative to SE events could suggest an enrichment for splicing changes associated with functional remodeling. For loss edges, we observed a positive correlation, with each SE event potentially contributing to multiple edge losses. This cascading effect was evident in extreme cases such as LAML patient TCGA-AB-2811, whose approximately 14,000 loss edges reflected extensive remodeling of key network hubs including KRAS (loss degree = 296), HSP90AA1 (loss degree = 234), HRAS (loss degree = 234), RAC1 (loss degree = 197), and ELAVL1 (loss degree = 194). For gain edges, most cancer types showed fewer gains relative to SE events, with brain cancers again standing out as having roughly equal or greater numbers of gain edges relative to their SE burden.

**Figure 3.**
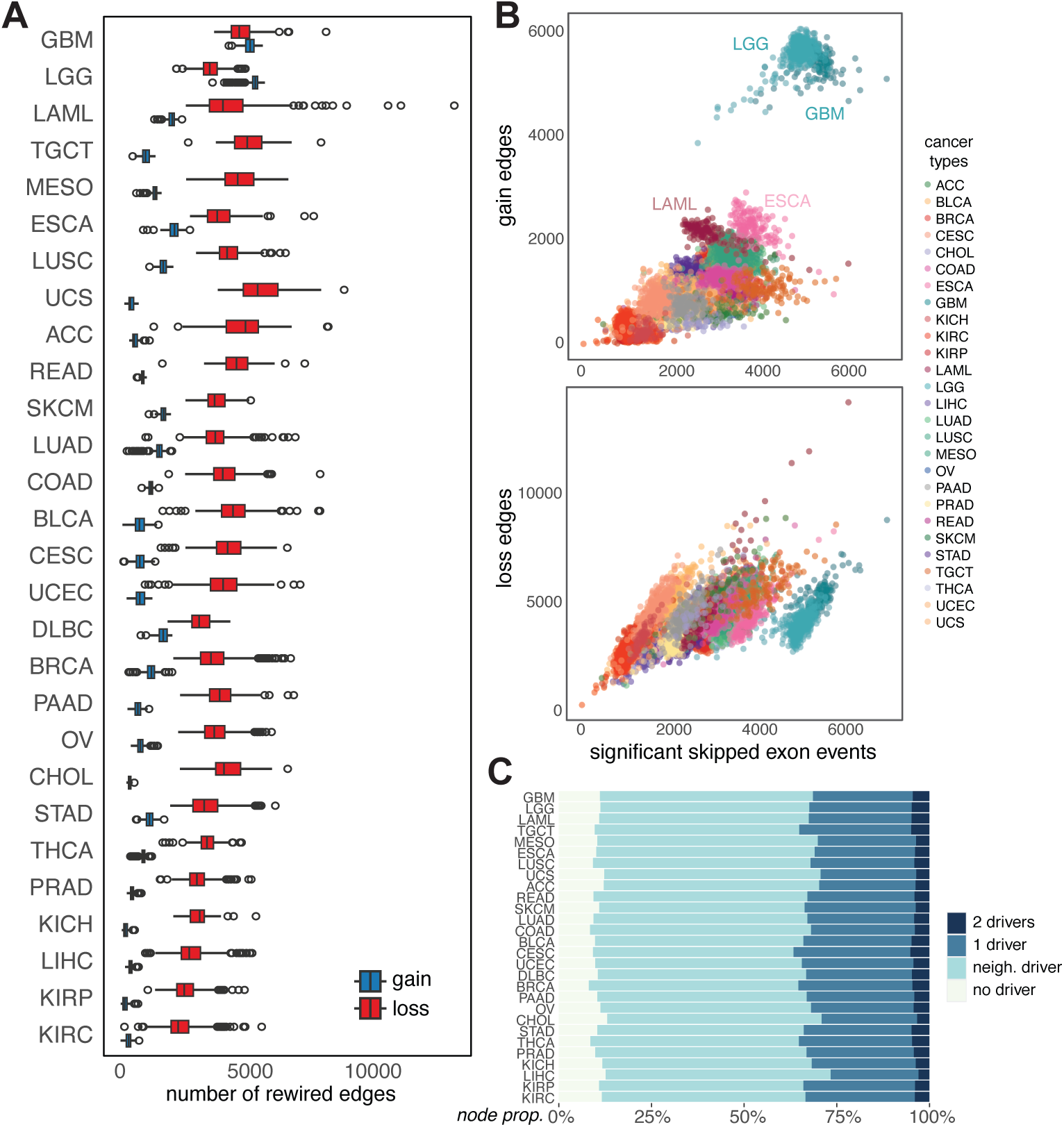
Splicing-induced rewiring of PPI networks across cancers. (A) Box plots summarize the distributions of gained (blue) and lost (red) edges for each cancer type, ordered by median of total number of edges in the largest connected component. The brain cancers GBM and LGG show a higher proportion of gains to losses. (B) Scatter plots show the relationship between the number of significant SE events (x-axis) and the number of rewired edges (y-axis) for individual tumor samples, separated by PPI rewired edge type (top gain edges, bottom loss edges). (C) Stacked bar plots for each cancer show the proportion of edges involving known cancer driver genes, from no involvement to both nodes in the edge being cancer driver genes.

We next investigated whether these rewiring events preferentially affect known cancer driver genes. Using driver gene annotations from the COSMIC census [24] and patient-level mutation calls, we found that driver genes consistently had significantly higher average degrees in the rewired networks than non-driver genes, regardless of whether the driver gene was mutated in that particular patient. Rank-biserial tests confirmed a moderate effect for driver versus non-driver status (rank-biserial = 0.24) and a negligible effect for mutated versus non-mutated genes (rank-biserial = 0.05; Table S2). The vast majority of rewired edges involved at least one driver gene either directly or through an immediate neighbor. On average, 28% of edges directly involved one driver, 5% involved two drivers, and a further 57% involved a neighbor of a driver, leaving only 10% of edges with no driver involvement (Figure 3C). This distribution was relatively consistent across cancer types, with some level of minor variation. CHOL, LIHC, adrenal cortical carcinoma (ACC), and UCS showed slightly higher proportion of driver-free edges (12-13%), while BRCA, COAD, THCA, and CESC showed modestly increased driver involvement (8-9% without drivers). The observation that driver genes are preferentially affected by splicing-driven rewiring regardless of their mutation status suggests that alternative splicing may remodel established oncogenic pathways through a mechanism that is complementary to, rather than redundant with, somatic mutations.

### Consensus network analysis identifies conserved rewiring programs and cancer type-specific hub genes

To move from individual patient networks to cancer type-level patterns, we constructed “consensus networks” summarizing the fraction of rewired edges retained across patients within each cancer type. For each cancer type and edge direction (gain or loss), we generated consensus networks at five conservation thresholds (0%, 25%, 50%, 75%, and 100%), where a threshold of 25%, for example, retains only edges present in at least 25% of patient networks for that cancer type. Comparing edge compositions across thresholds revealed highly heterogeneous patient-specific rewiring: few edges were conserved across all patients within a cancer type, and even fewer appeared across multiple cancer types (Figure 4A). Nonetheless, for most cancer types, a notable subset of edges appeared in at least one quarter of patient samples, suggesting the existence of potential molecular subtypes with shared rewiring signatures (Figure S2). Conservation patterns also differed by edge direction: gain edges showed lower within-and across-cancer conservation than loss edges, suggesting that core cancer processes may be more strongly reflected in interaction losses.

**Figure 4.**
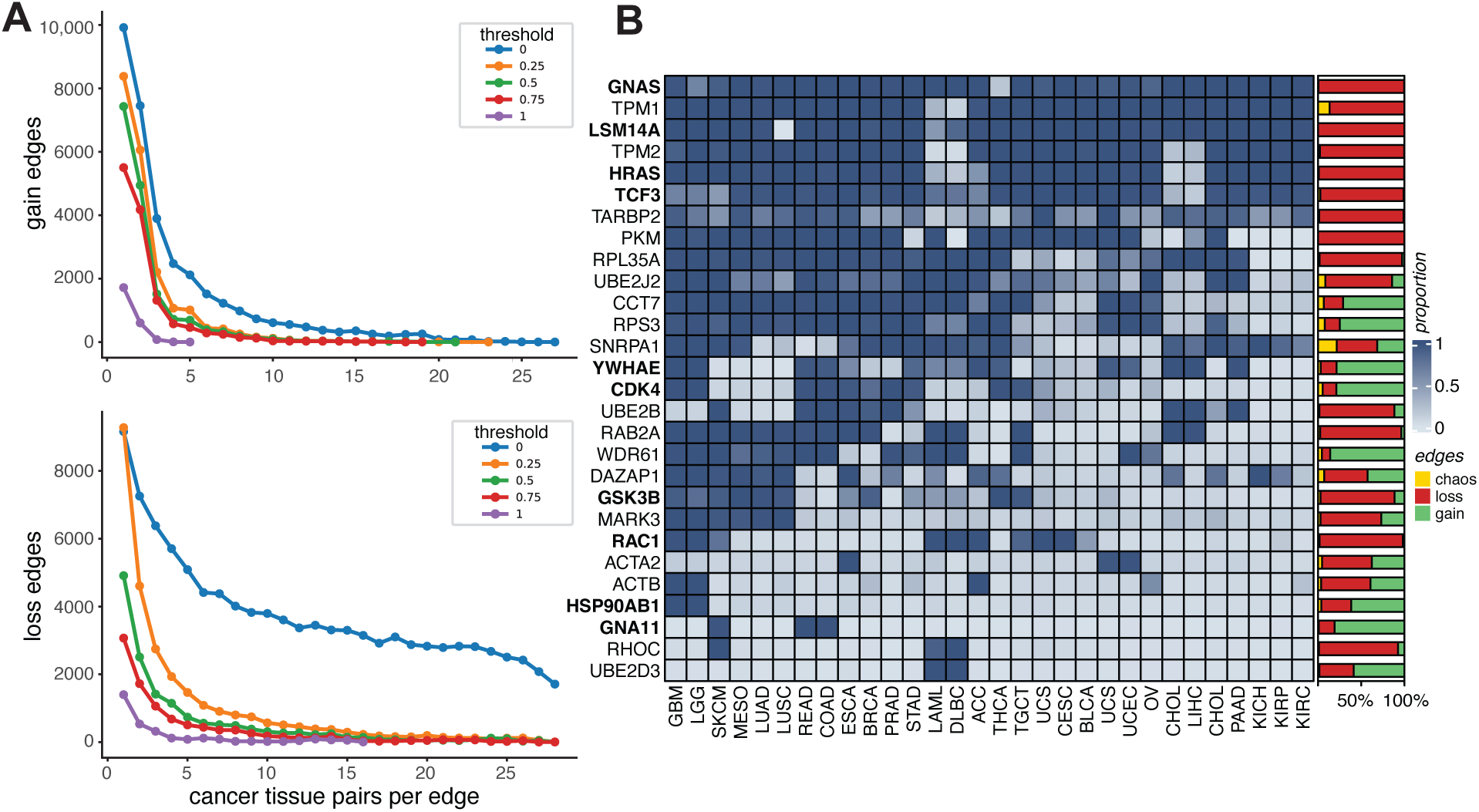
Node and edge conservation across cancers. (A) Frequency distribution of edges filtered by internal consistency within a cancer type (0, 0.25, 0.5, 0.75, 1). For each threshold, only edges that occur in that minimum proportion of samples for a given cancer type are retained. The x-axis shows the number of cancers in which an edge is observed and the y-axis the total edge count, subset by either gain (top) or loss edges (bottom) for a given threshold. (B) Heatmap of the top rewired gene for each cancer type (selected without duplicates based on rank). Here, genes are ordered by the average ratio of rewired to background degree, sample presence, and total degree. Bold names indicate known cancer drivers. Heatmap colors represent the average ratio of rewired gene degree to background degree for each cancer type, with an adjacent bar chart showing the relative proportions of rewired edge types across all samples.

We next examined how the biological functions of rewired edges vary with their conservation patterns. Edges that were highly conserved within a cancer type but unique across cancer types (observed in ≤ 2 cancers at threshold = 1; 1,931 loss and 2,321 gain edges) showed distinct functional signatures. Loss edges in this set were strongly enriched for signaling-related processes, including small GTPase-mediated signal transduction (*padj* = 6.16*e*^−61^), Ras protein signal transduction (*padj* = 5.72*e*^−34^), and protein autophosphorylation (*padj* = 1.40*e*^−19^; Table S5). In contrast, gain edges were enriched for splicing-related functions, including modification-dependent protein catabolic process (*padj* = 1.01*e*^−25^), RNA processing (*padj* = 5.47*e*^−18^), and RNA splicing via transesterification reactions (*padj* = 1.10*e*^−17^; Table S5), suggesting potential splicing feedback mechanisms.

We then examined edges that were conserved across many cancer types (observed in ≥ 10 cancers at thresholds ≥ 0.75; 1,273 loss and 169 gain edges), which may represent core splicing signals shared across tumors regardless of tissue of origin. Loss edges in this pan-cancer set were again enriched for GTPase-related signaling terms, reinforcing the pattern observed in cancer-specific edges. Gain edges, however, were enriched for a distinct set of processes centered on SNARE-mediated membrane fusion and vesicle trafficking (e.g., membrane fusion, *padj* = 3.39*e*^−14^), as well as cytoplasmic translation (*padj* = 3.78*e*^−9^) and translational initiation (*padj* = 4.47*e*^−7^) (Table S4). Comparing the gene sets underlying these edge groups, loss-edge genes showed substantial overlap between the cancer-specific and pan-cancer sets (Jaccard index = 0.49, Table S3), suggesting that tumor type-specific and cross-cancer interaction losses converge on similar core pathways. In contrast, gain-edge genes were largely distinct between the two sets (Jaccard index = 0.08), reflecting greater heterogeneity in the processes affected by gained interactions.

To more precisely characterize the conservation of gene-level remodeling across cancer types, we identified the most rewired gene for each cancer type after normalizing for background PPI network degree (see Methods). The resulting set of 28 genes spans the full range from pan-cancer to cancer type-specific rewiring (Figure 4B, Table S6). The most broadly conserved members were almost uniformly loss-dominated. Six genes exhibited consistent loss-type rewiring across nearly all tumor types: GNAS (99.0% of edges), HRAS (98.7%), LSM14A (99.9%), TCF3 (96.3%), TPM1 (86.6%), and TPM2 (97.6%). Four of these (GNAS, HRAS, LSM14A, TCF3) are known oncogenes [24], and the remaining two (TPM1 and TPM2) have well-documented roles in splicing and isoform regulation [25]. Only two genes with similar pan-cancer distribution showed widespread gain-type rewiring: CCT7 (70.9% of edges) and RPS3 (74.8%).

Whereas broadly conserved genes were predominantly loss-rewired, genes with more cancer type-restricted remodeling tended toward heterogeneous mixtures of gain and loss, and several were strongly gain-enriched. WDR61 had the highest proportion of gain-edge rewiring among top-ranking genes (86.9%). CDK4, a frequently altered driver, exhibited gain-dominant rewiring (78.8%) across a subset of solid tumors, including those of the brain, testis, thyroid, esophagus, breast, prostate, and stomach. HSP90AB1 displayed primarily gain rewiring (61.6%) restricted to brain tumors, consistent with its known alteration frequency in GBM [26]. GNA11, a well-known driver in uveal melanoma (not included in our subset of cancer types), also showed predominantly gain rewiring (80.9%), largely restricted to colorectal and skin cancers. Although no completed studies have firmly established a role for GNA11 in colorectal cancer, one clinical trial is currently investigating this possibility (NCT03947385), and prior work has implicated it in multiple skin malignancies [27].

Aside from directional polarity, three additional genes stood out. SNRPA1, a core spliceosomal component [28] and a previously reported prognostic marker in breast cancer metastasis [29], carried the highest proportion of chaos edges (21.6%) among top-ranking genes, suggesting substantial and directionally discordant disruption at a central regulatory node. Meanwhile, two non-driver genes, RHOC and UBE2D3, appeared to be highly specific to hematologic cancers, with limited signal elsewhere. The contrast between broadly conserved loss-type rewiring at signaling hubs and more heterogeneous, gain-enriched rewiring at cancer type-specific genes suggests that these could represent distinct modes of splicing-driven network remodeling.

### Bidirectional switches and non-polarized edges reveal distinct modes of network rewiring

To evaluate how consistently edges are rewired across cancer types, we defined a consistency metric as the difference between the number of gain and loss events for an edge, normalized by the total number of samples for each cancer type. Values near –1 indicate that loss events overwhelmingly dominate, whereas values near +1 indicate predominant gain rewiring for a specific edge. Restricting to edges present in more than 50% of samples within a given cancer type, we found that directionality is typically conserved, with consistency scores clustered near the extremes of-1 or +1 (Figure S3). Across cancer types, edges skewed toward loss-dominated rewiring (median consistency: –0.65) consistent with the pan-cancer asymmetry observed earlier. Brain cancers were again the exception, with a strong shift toward gain-dominated rewiring (median consistency: 0.86), reinforcing the pattern that gain rewiring is a hallmark of these tumors.

Against this broad pattern of within-cancer directional coherence, a subset of edges displayed divergent behavior across cancer types, with strong gains in some cancers and strong losses in others (Figure 5A). We defined these bidirectional edges as those appearing as gains in at least 70% of samples from three or more cancer types and as losses in at least 70% of samples from three other cancer types. A total of 221 edges met these criteria, with four genes accounting for the majority of bidirectional interactions: CDK7 (23.1% of edges), FN1 (14.5%), PTPN6 (13.6%), and SNRPA1 (13.1%). The biology of these four hubs supports a switch-like role in cancer. CDK7 is a cyclin-dependent kinase that also phosphorylates spliceosomal factors such as SF3B1 [30] and has been emerging as a promising target for selective inhibition in clinical trials [31]. FN1 has been found to undergo cancer-associated alternative splicing that can play opposing roles in thyroid cancer [32], and in general has been controversial for its role in cancer, promoting tumor growth in some cancers (e.g., colorectal [33], breast [34]), while suppressing it in others (e.g., melanoma [35]). PTPN6 encodes a tyrosine phosphatase that is expressed primarily in hematopoietic cells via a tissue-specific promoter [36], whose aberrant splicing has been implicated in acute myeloid leukemia [37]. SNRPA1 is, interestingly, the core spliceosomal component gene we previously identified as having the highest chaos edge count among top pan-cancer rewired genes (Figure 4B), which has also been identified as a mediator of metastatic breast cancer [29]. Beyond these four dominant hubs, ten additional genes occurred two or more times among bidirectional edge-partner sets. Two, CASP9 and GSK3B, are annotated as drivers in the COSMIC Cancer Gene Census [24]. Others have been linked to cancer-associated regulatory programs, including HNRNPH1 and DAZAP1, which were implicated in disrupted RNA-binding and splicing regulation in mantle cell lymphoma [38], and SRPK2, which has been reported to influence lipogenesis and cancer cell growth [39]. Together, these bidirectional rewiring patterns reveal genes that may function as context-dependent switches whose rewiring direction varies systematically across cancer types, reflecting splicing programs that selectively reconfigure shared regulatory nodes to support tissue-specific oncogenic states.

**Figure 5.**
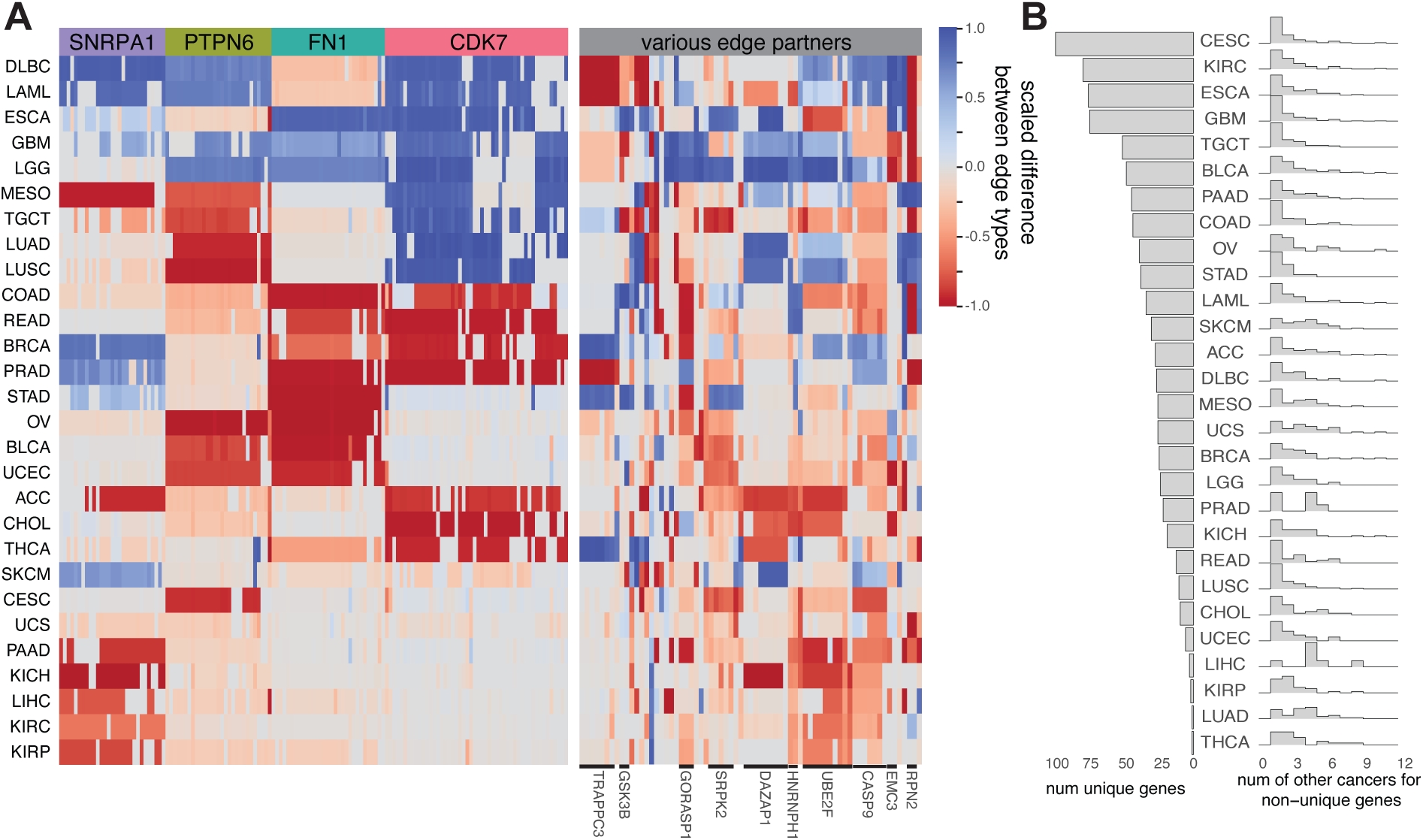
Recurring polarized and non-polarized rewiring patterns. (A) Heatmap of consistency values (the proportion of edges with a conserved directionality) across cancer types. Values colored in blue represent more conserved gain edges, where those in red highlight conserved loss edges. Top repeating members of these edges are shown in colored column bands, while the right column (in gray) represents the remaining edges. Genes occurring two or more times within the various edge-partner sets are highlighted and reported at the bottom of the plot. (B) Composition of the non-polarized edges across cancer types ranked by the number of unique genes within these edges. The left side of the figure shows a bar plot of the counts of the unique cancer-specific genes in the non-polarized edge set. The histogram on the right tallies up how many of the non-unique genes are shared across other cancer types.

Complementing these bidirectional patterns, a second class of edges showed consistent presence within a cancer type but no directional bias, suggesting splicing changes that mix gain and loss events in roughly equal proportion across patients. We isolated edges with consistency values between-0.25 and 0.25 that were present in at least 50% of samples for a given cancer type, yielding 2,163 unique edges spanning 1,501 genes. GO enrichment analysis revealed significant associations with core tumorigenic processes, including cell division (*padj* = 2.36*e*^−11^), regulation of transferase activity (*padj* = 1.78*e*^−9^), and chromosome organization (*padj* = 4.95*e*^−9^). The extent of non-polarized rewiring varied substantially across cancer types, ranging from 217 edges (across 234 genes) in GBM to 7 edges (across 10 genes) in LIHC. The genes underlying these edges were largely cancer type-specific, regardless of edge directionality (mean Jaccard index = 0.03; Figure 5B). CESC had the most unique gene set, enriched for fibroblast signaling (*padj* = 1.03*e*^−11^), while THCA had the least number of unique genes, with shared genes enriched for protein post-translational modification processes (*padj* = 5.77*e*^−4^, Table S7). GBM and LUSC had the highest overlap in genes (overall Jaccard = 0.24), particularly for loss-biased edges (Jaccard = 0.31), while DLBC and SKCM showed the greatest similarity for gain-biased edges (Jaccard = 0.27). Among all genes in this non-polarized set, Ubiquitin C (UBC; mean consistency =-0.05) appeared in the greatest number of cancer types (11 in total), suggesting a widely shared but variable rewiring pattern across patients, which may have meaningful clinical implications [40].

### Outlier rewiring predicts survival independent of stage and mutational burden

To assess whether patient-level rewiring carries prognostic information, we investigated whether tumors whose networks deviate from the consensus for their cancer type exhibit distinct clinical outcomes. We applied DBSCAN to identify outlier samples within each cancer type, using two features per sample: the number of significant SE events and the number of edges in the largest connected component of the rewired network (Methods). Gain and loss edges were analyzed separately, yielding 134 gain outliers and 74 loss outliers, corresponding to an average of 2-5 outliers per edge type and cancer type (Figure 6A).

**Figure 6.**
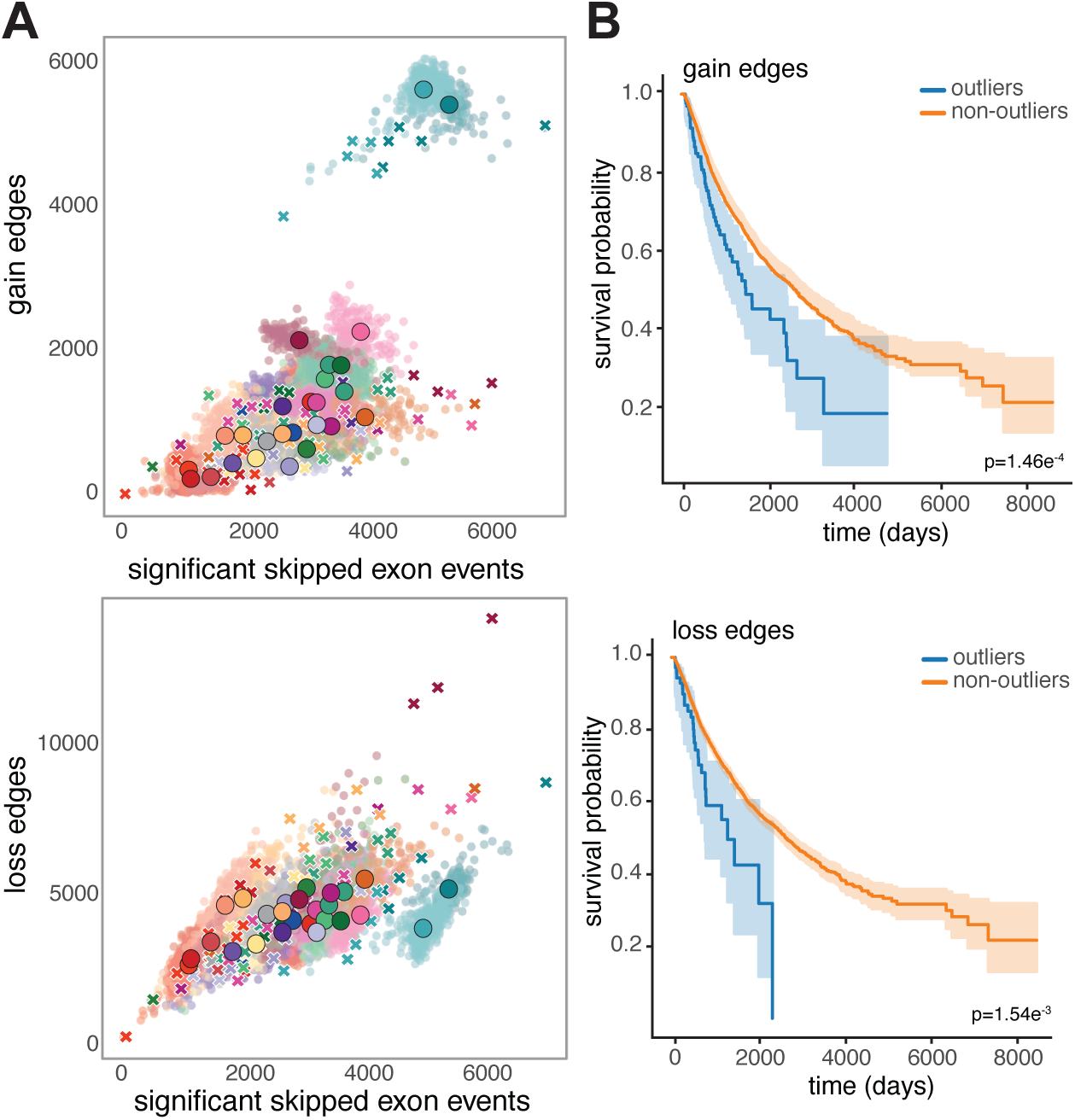
Patients with outlier rewired networks have distinctly worse survival. (A) Scatter plot showing the relationship between the number of significant SE events versus the number of gain (top) or loss (bottom) edges in the largest connected component of the rewired network. Individual patient samples are represented by transparent dots, centroids for each cancer types are shown as a large solid dot and outlier patient samples are marked by ‘X’s. (B) Kaplan–Meier survival curves comparing overall survival between outlier (blue) and non-outlier (orange) patients across all cancer types. Shaded regions depict 95% confidence intervals.

Outlier patients had significantly worse overall survival than non-outliers for both edge types (log rank test, gain: *p* = 1.46*e*^−4^; loss: *p* = 1.54*e*^−3^; Figure 6B). To analyze whether this association was independent of established clinical features, we fit a multivariate Cox proportional hazards model stratified by cancer type with tumor stage, gender, age at diagnosis, and outlier status as covariates (Figure S4). Gain edge outliers showed a 56% increased risk of death compared to non-outliers (HR = 1.56, *p* = 6.99*e*^−3^), a hazard ratio second only to that of late-stage disease (Stage III, IV). Loss edge outliers showed a comparable point estimate but did not reach statistical significance (HR = 1.47, 95% CI: 0.95–2.28; *p* = 0.085), and the significance of this effect was sensitive to DBSCAN parameter choice (Methods). These results suggest that loss outliers may largely reflect features already captured by clinical staging, whereas gain outliers contribute prognostic information that is complementary to and independent of standard clinical covariates.

To test whether outlier status reflects features of the somatic mutation landscape, we compared mutation profiles between outliers and non-outliers for each edge type. Overall mutation count did not differ significantly based on outlier status (Wilcoxon rank sum test, gain: *p* = 0.56; loss *p* = 0.89). We also examined the composition of mutation types based on outlier status (Methods). Gain outliers showed modest compositional shifts relative to non-outliers (MANOVA *p* = 0.001), but the effect size was negligible (Wilks’ lambda=0.9965) and the signal was concentrated in rare mutation classes (collectively under 2.5% of mutations), with no major class differing significantly after correction (Table S8). For loss outliers, mutation type composition was not significantly different from non-outliers (MANOVA *p* = 0.21, Wilks’ lambda=0.9978). Thus, we did not find evidence that outlier status for either edge type was meaningfully driven by either mutational load or mutational composition.

## Discussion

This work establishes splicing-driven PPI network rewiring as a structured, directionally asymmetric, and prognostically informative dimension of cancer biology. A central theme that emerges is that splicing relates to somatic mutation in distinct ways: in some respects it converges on the same oncogenic circuitry that mutation targets, and in others it carries signal that mutation does not. Driver genes are preferentially rewired across patient networks regardless of whether they are mutated in a given patient, leading us to hypothesize that splicing is as an additional route to altering the function of established cancer drivers. The loss-dominated rewiring conserved across cancer types likewise converges on Ras and GTPase signaling, pathways that are themselves frequent targets of oncogenic somatic alteration. We see this most clearly in the prognostic analysis where tumors whose networks deviate most from their cancer-type consensus, particularly those dominated by interaction gains, show significantly worse survival that is not accounted for by mutational burden, mutational composition, or tumor stage. Splicing-driven remodeling thus neither simply recapitulates somatic alteration nor acts wholly apart from it, intersecting with mutation on shared oncogenic pathways while contributing prognostic information of its own.

Across cancer types, brain cancers stood out as a major exception to the loss-dominated rewiring pattern in most cancers, where both LGG and GBM exhibited gain-dominated rewiring. We speculate that this divergence may reflect the inherently complex splicing landscape of neural tissues, where healthy brain tissue exhibits the highest level of alternative splicing among all human tissues [41–43]. This pattern is particularly pronounced in the cerebellum, which shows a high rate of novel splicing events and significant enrichment for RNA-binding proteins, which are known to regulate splicing [42]. The elevated interaction gains observed here in brain cancers could reflect an adaptive mechanism or a consequence of the unique neural environment. The finding takes on additional interest in light of our prognostic analysis, where gain-dominated rewiring is associated with worse survival across cancer types. The fact that brain cancers as a whole exhibit gain-dominated networks raises the possibility that gain-dominated rewiring outliers in non-brain cancers could reflect activation of neural-like splicing programs.

Our analysis identified four major genes (CDK7, FN1, PTPN6, SNRPA1) that function as bidirectional switches, undergoing strong gain rewiring in some cancer types and strong loss rewiring in others. Each of these genes has independent literature support for cancer-associated splicing changes, but no prior case-by-case study has unified them or revealed their cross-cancer switch behavior. This pattern emerged because we were able to perform directional inference at the network level, which distinguishes our approach from prior PPI rewiring methods that treat splicing-induced changes as undirected disruptions. The four span distinct mechanistic categories, from a transcription and splicing kinase (CDK7) to a structural ECM gene with well-documented isoform switching (FN1), as well as a tissue-restricted signaling phosphatase (PTPN6) and a core spliceosomal regulator (SNRPA1). Thus, bidirectional switching seems to be a property of multiple gene classes rather than a single regulatory mechanism. The presence of clinically actionable targets in this set, including CDK7 with multiple inhibitors in clinical trials [31], suggests that pan-cancer rewiring patterns may help identify shared therapeutic vulnerabilities across cancers with otherwise distinct tissue origins.

Among the switch genes, SNRPA1 had the highest proportion of chaos edges of any top-ranked gene, reflecting splicing changes that simultaneously perturb domain-domain interactions in opposing directions. This contrasts with the majority of rewired edges in our dataset, which arise from more consistent exon exclusion or inclusion events. The presence of such discordant, multidirectional remodeling suggests that SNRPA1 may influence protein interaction landscapes in a particularly destabilizing manner, potentially capturing the widespread transcriptomic and phenotypic plasticity attributed to its splicing activity. Furthermore, it points to the possibility that chaos edges, rather than representing analytical noise, may carry biological signal at genes whose splicing activity destabilizes the interaction landscape itself.

We acknowledge several limitations in the interpretation of these findings. Our analysis predicts interaction-level changes from exon-level splicing data and protein domain annotations, but the predictions are computational and would benefit from orthogonal validation by isoform-resolved interaction assays. While such datasets are still limited, there are emerging efforts to systematically build these resources [9, 44]. Tissue matching between TCGA tumors and GTEx normals is also imperfect for some cancer types, particularly LIHC, CHOL, MESO, and LUSC, where the divergence between cancers sharing a tissue background may partly reflect suboptimal reference matching rather than purely biological differences. We have focused exclusively on exon skipping events to capture the most prevalent and mechanistically interpretable class of alternative splicing, but other splicing event types (intron retention, alternative splice sites, mutually exclusive exons) may also contribute additional rewiring not captured here.

Looking ahead, further integration of this analysis paradigm with complementary data, including gene expression, DNA methylation, and copy number variation, will help elucidate the dynamic interplay underlying interaction rewiring across cancer types, and the link between gain-dominated rewiring outliers and worse survival merits further investigation as a candidate prognostic marker. More broadly, the patterns identified here establish splicing-driven interaction rewiring as a distinct and prognostically relevant axis of oncogenic network remodeling, surfacing specific interactions that may mediate tumor aggressiveness and warrant deeper mechanistic and clinical study.

## Methods

### Data preprocessing

#### Protein-protein and domain-domain interactions

Protein-protein interactions (PPIs) and domain-domain interactions (DDIs) were processed as previously described [15]. Briefly, human PPIs were aggregated from 8 databases (BioGRID [45], DIP [46], HIPPIE [47], HPRD [48], Human Interactome [49], IntAct [50], iRefIndex [51], MIPS [52]), and DDIs were obtained from 4 sources (3did [53], DOMINE [54], IDDI [55], iPFAM [56]). For predicted DDIs, only interactions with a confidence score above 0.5 were included. Protein domain locations were translated to genomic coordinates using the Ensembl BioMart API and the biomaRt R package [57] and indexed with Tabix [58] for fast lookup. The final scaffold consensus network has 20,286 nodes and 793,078 edges in the consensus PPI and 17,450 domain pairs in the DDI set.

#### Splicing data

Skipped exon (SE) splicing quantifications for 7,950 TCGA tumor samples spanning 28 cancer types [17] and 5,968 GTEx normal tissue samples [18] were obtained from the IRIS database [20], which uses rMATS [59] to compute percent spliced in (PSI or *ψ*) values. The specific exons detected in the TCGA sample and GTEx normal tissue samples are not necessarily identical, and we only focus on the exons captured in both. For each tumor sample, we computed the change in Δ*ψ_i_*for an exon *i* relative to the mean of its matched normal tissue background:

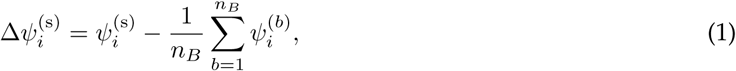

where 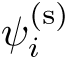 is the PSI for exon *i* in a TCGA cancer sample, 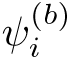 is the PSI for exon *i* in background GTEx normal sample *b*, and *n_B_* is the total number of GTEx background samples. We assign an empirical p-value for 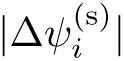 by constructing an empirical cumulative distribution function (CDF) for each exon *i* based on the background GTEx samples (see [15] for details). Exons were considered significantly altered if both 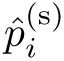 *<* 0.05 and |Δ*ψ*| *>* 0.05 as in the original Splitpea study.

#### Tissue matching

Each TCGA cancer type was manually mapped to its closest GTEx normal tissue (Table S1). Cancer types without a clear tissue counterpart in IRIS, such as sarcoma, were excluded. For two cancer types, we identified two plausible tissue matches and used both as background, adjusting sample sizes to ensure balanced representation. Specifically, for uterine carcinosarcoma (UCS), we combined Cervix Uteri (11 samples) and Uterus (95 samples) by oversampling Cervix Uteri to reach approximately one-third the Uterus sample count and randomly subsampled Uterus down to 32 samples. For cholangiocarcinoma (CHOL), Liver (146 samples) and Small Intestine (111 samples) were more similar in size, so we randomly subsampled Liver to match the 111 samples of Small Intestine without any oversampling.

#### Somatic mutation data

Sample-level somatic mutation calls for TCGA tumors were obtained from the TCGA MC3 (Multi-Center Mutation Calling in Multiple Cancers) dataset [60] using the TCGAmutations resource (available at: https://github.com/PoisonAlien/TCGAmutations). Mutation annotation and processing were performed using the maftools R package [61]. We included the following mutation classifications in our analyses: Missense Mutation, Nonsense Mutation, Frame Shift Del, Frame Shift Ins, Splice Site, In Frame Del, In Frame Ins, and Nonstop Mutation. Per-sample mutation counts were computed as the total number of qualifying mutations, and per-gene mutation status was defined as the presence of at least one qualifying somatic mutation. Cancer driver gene annotations were obtained from the COSMIC Cancer Gene Census [24].

#### Generating patient-specific rewired PPI networks

We used Splitpea [15] with its default parameters and recommended pipeline. Splitpea integrates the significant SE events with consensus PPI and DDI networks to identify protein interactions whose underlying domain-mediated contacts are altered by splicing changes. For each interacting protein pair, Splitpea evaluates whether the constituent domain-domain interactions are consistently increased (gain edge), consistently decreased (loss edge), or affected in opposing directions (chaos edge), and reports both an edge weight derived from the Δ*ψ_i_* values and a corresponding directionality label. Running Splitpea for each of the tumor samples yielded 7,950 patient-specific rewired networks.

#### Consensus networks

To identify rewiring events conserved across patients within each cancer type, we constructed consensus networks by filtering edges based on their frequency across patient-specific networks for that cancer type. For each cancer type, gain edges and loss edges were processed separately; chaos edges were excluded due to their ambiguous directionality. We evaluated five consensus thresholds: 0, 0.25, 0.5, 0.75, and 1. At each threshold *t*, an edge is retained only if it appears in at least *t* fraction of patient samples for that cancer type. For *t* = 0, the edge needs to appear in at least one sample. This resulted in a total of 10 consensus networks per cancer type (five thresholds x two directions). All consensus networks were constructed using the NetworkX package [62].

#### Top rewired genes ranking

To identify the most rewired genes for each cancer type, we ranked genes using three metrics: normalized degree proportion, gene presence, and overall degree. The normalized degree proportion (NDP) for gene *g* in cancer type *c* is defined as

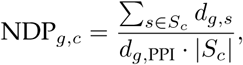

where *S_c_* is the set of samples for cancer type *c*, *d_g,s_* is the observed degree of gene *g* in sample *s*, and *d_g,_*_PPI_ is the degree of *g* in the unmodified background PPI network (its maximum possible degree); a ratio of 1 indicates that the gene is fully rewired, as its observed connections equal the maximum possible. Ties in normalized degree proportion are then broken by gene presence followed by overall degree. Gene presence is defined by the number of samples in which the gene appears, while overall degree is the total number of connections over all the patient networks summed for that cancer. To produce a non-redundant set of top-ranked genes across cancer types for visualization (Figure 4B), we iterated through cancer types in descending cohort size and, for each, selected the highest ranked gene that had not yet been assigned to another cancer type. The heatmap is clustered and visualized using the default parameters from the ComplexHeatmap package [63].

#### Consistency analysis

To characterize the directional consistency of rewiring events within and across cancer types, we defined a per-edge, per-cancer consistency metric. For each edge *e* in cancer type *c*:

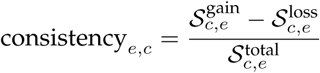

where S_gain_ is the number of samples in which the edge is labeled as gained, S_loss_ is the number of samples in which the edge is labeled as loss, and S_total_ is the total number of samples with observed significant skipped exon events for that edge in the given cancer type. The metric ranges from-1 (all observations are losses) to +1 (all observations are gains), with values near 0 indicating that gain and loss events occur in approximately equal proportions across patients. We refer to edges with consistency values close to either extreme as *directionally polarized* and those with values near 0 as *non-polarized*. Note that consistency here refers to within-cancer directional agreement, not simply whether the edge is consistently observed across samples.

#### Bidirectional switch edges

To identify edges whose rewiring direction varies systematically across cancer types, we defined bidirectional switches as edges meeting two criteria: (i) consistency ≥ 0 in at least 70% of samples in 3 cancer types, and (ii) consistency ≤ 0 in at least 70% of samples in 3 other cancer types.

#### Non-polarized edges

To identify edges with mixed gain and loss rewiring within cancer types without a strong directional polarization, we selected edges with consistency values between-0.25 and 0.25 that were present in at least 50% of samples of a given cancer type.

### Outlier survival analysis

#### Outlier identification

For each cancer type, we identified patient samples whose rewired networks deviated substantially from the cancer type consensus by applying DBSCAN [64] to two per-sample features: the number of significant skipped exon events and the number of edges in the largest connected component of the rewired network. We restricted to the largest connected component to focus on the primary rewired subnetwork in each sample, which captures the bulk of rewiring activity while excluding small disconnected fragments that can contribute disproportionate variance for samples with sparse rewiring. Gain edges and loss edges were analyzed separately, producing two outlier sets per cancer type. Features were standardized within each cancer type using z-score normalization, and DBSCAN was applied with parameters eps=0.7 and min_samples=2 using the scikit-learn [65] implementation.

#### Survival modeling

Clinical metadata (survival time, event status, tumor stage, age at diagnosis, gender) were obtained from the Genomic Data Commons Data Portal [66]. Kaplan–Meier survival curves and log-rank tests were computed using the lifelines Python package [67]. To assess whether outlier status carries prognostic information independent of established clinical features, we fit multivariate Cox proportional hazards models using the statsmodels Python package [68], stratified by cancer type and adjusted for tumor stage, age at diagnosis, and gender. Gain edge outliers identified by DBSCAN with min_samples below 4 remained consistently significant in both Kaplan-Meier curves and the Cox model. Loss edge outliers were significant when min_samples was below 4 in the Kaplan-Meier analysis, but their significance in the Cox model was sensitive to the choice of the eps parameter.

#### Somatic mutation composition comparison

For each patient, we computed the proportion of somatic mutations in each mutation class, analyzing gain and loss outliers separately. Mutation classes with <2% of all mutations (Frame Shift Ins, In Frame Del, In Frame Ins, Nonstop Mutation, Translation Start Site) were aggregated into an “Other” category to avoid instability from sparse proportions. For each of the classes (Frame Shift Del, Missense Mutation, Nonsense Mutation, Splice Site, Other), we compared composition between outliers and non-outliers using a Wilcoxon rank-sum test on per-patient proportions with Benjamini-Hochberg for multiple hypothesis test correction. Because mutation class proportions are compositional, we also applied an additive log-ratio (ALR) transform to each patient’s mutation composition and tested whether outlier status was associated with overall composition using the MANOVA model that also adjusts for overall mutational burden and cancer type: ALR(mutation composition) ∼ outlier status + log(mutation count) + *C*(cancer type).

### Gene Ontology enrichment analysis

To investigate the biological processes associated with network alterations, we performed Gene Ontology (GO) enrichment analysis in R using the clusterProfiler package [69] with annotations from the org.Hs.eg.db database [70]. For each set of genes we first mapped to Entrez IDs, and unmapped genes were discarded. Enrichment was tested against a background gene set consisting of all genes present in the unmodified consensus PPI network. Benjamini–Hochberg was used for multiple hypothesis test correction.

## Data and code availability

All analysis code is available on GitHub at https://github.com/ylaboratory/ splitpea-pancancer-analysis, released under the BSD 3-clause license for open source use, and generated networks are available for download on Zenodo at https://zenodo.org/records/20767005 under the CC BY 4.0 license.

## Supplementary Materials

**Table S1.** TCGA cancer type matched with GTEx tissue in the Splitpea analysis.

| TCGA | GTEx Match |
| --- | --- |
| ACC | adrenal gland |
| BLCA | bladder |
| BRCA | breast |
| CESC | cervix uteri |
| CHOL | liver, small intestine |
| COAD | colon |
| DLBC | blood |
| ESCA | esophagus |
| GBM | brain |
| KICH | kidney |
| KIRC | kidney |
| KIRP | kidney |
| LAML | blood |
| LGG | brain |
| LIHC | liver |
| LUAD | lung |
| LUSC | lung |
| MESO | lung |
| OV | ovary |
| PAAD | pancreas |
| PRAD | prostate |
| READ | colon |
| SKCM | skin |
| STAD | stomach |
| TGCT | testis |
| THCA | thyroid |
| UCEC | uterus |
| UCS | uterus, cervix uteri |

**Table S2.** Average degrees for driver and non-driver genes stratified by mutation status.

| Category | Average degree (mean ± s.e.m.) |
| --- | --- |
| Mutated driver genes | 7.32 ± 0.109 |
| Non-mutated driver genes | 7.11 ± 0.014 |
| Mutated non-driver genes | 3.47 ± 0.019 |
| Non-mutated non-driver genes | 3.31 ± 0.002 |

**Table S3.** Jaccard similarity between loss and gain genes across conservation thresholds.

| Set 1 | Set 2 | Jaccard |
| --- | --- | --- |
| Loss (2 cancers, thresh 1) | Gain (2 cancers, thresh 1) | 0.0572 |
| Loss (2 cancers, thresh 1) | Loss (10 cancers, thresh 0.75) | 0.4939 |
| Loss (2 cancers, thresh 1) | Gain (10 cancers, thresh 0.75) | 0.0319 |
| Gain (2 cancers, thresh 1) | Loss (10 cancers, thresh 0.75) | 0.0743 |
| Gain (2 cancers, thresh 1) | Gain (10 cancers, thresh 0.75) | 0.0774 |
| Loss (10 cancers, thresh 0.75) | Gain (10 cancers, thresh 0.75) | 0.0514 |

**Table S4.** Top 20 enriched GO terms for genes in loss and gain edges in consensus networks at threshold *≥* 0.75 across *≥* 10 cancers.

**Table S5.** Top enriched GO terms for genes in loss and gain edges in consensus networks at threshold = 1 and observed in *≤* 2 cancers.

**Table S6.** Top 50 ranked genes for each cancer based on normalized degree proportion, gene presence, and overall degree.

**Table S7.** Enriched GO terms for non-polarized edges for each cancer type.

**Table S8.**
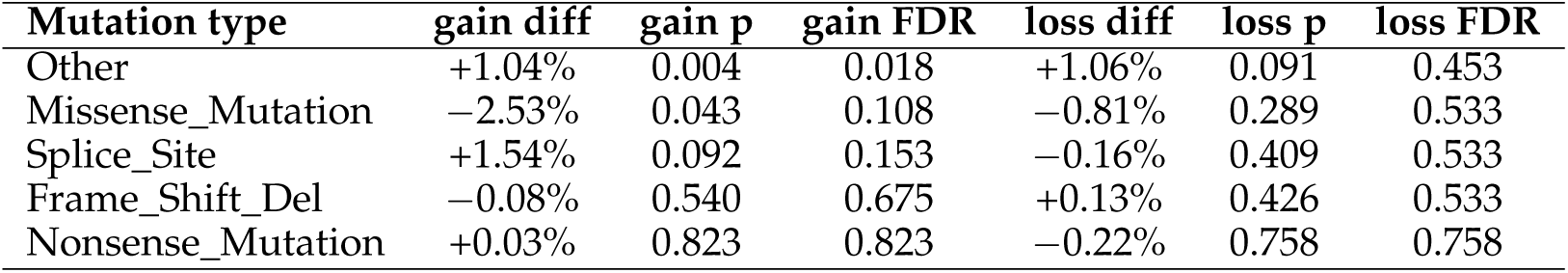
Wilcoxon rank-sum test comparing per-patient mutation-type fractions between outlier and non-outlier patients (gain outliers: n=134; loss outliers: n=74; total cohort: n=7,891); “diff” is the mean fraction difference (outlier minus non-outlier), with FDR correction applied via the Benjamini-Hochberg procedure; rare mutation types (<2% of total mutations) were collapsed into “Other.”

---

**Figure S1.**
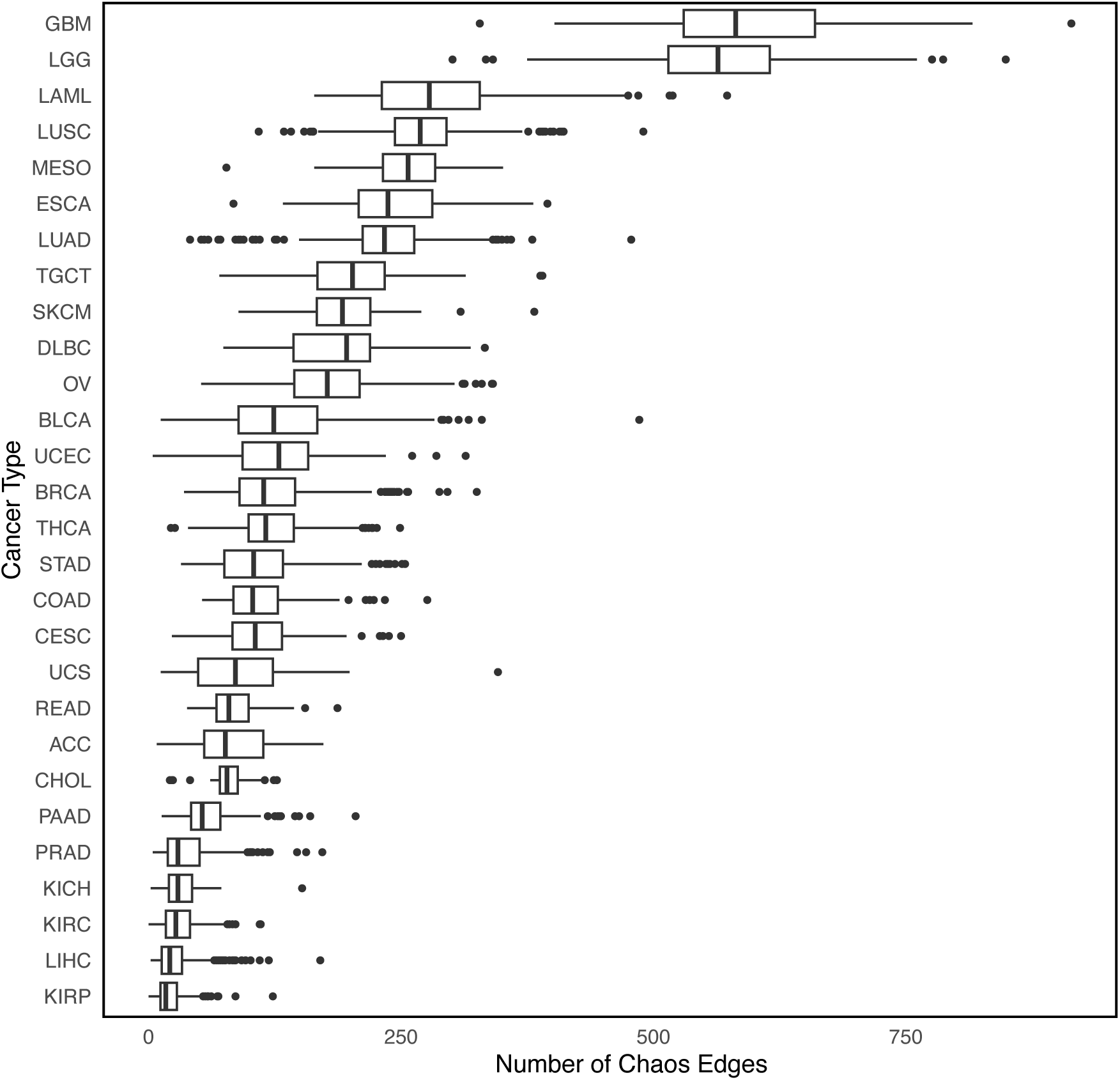
Box plots summarize the distributions of chaos edges for each cancer type.

**Figure S2.**
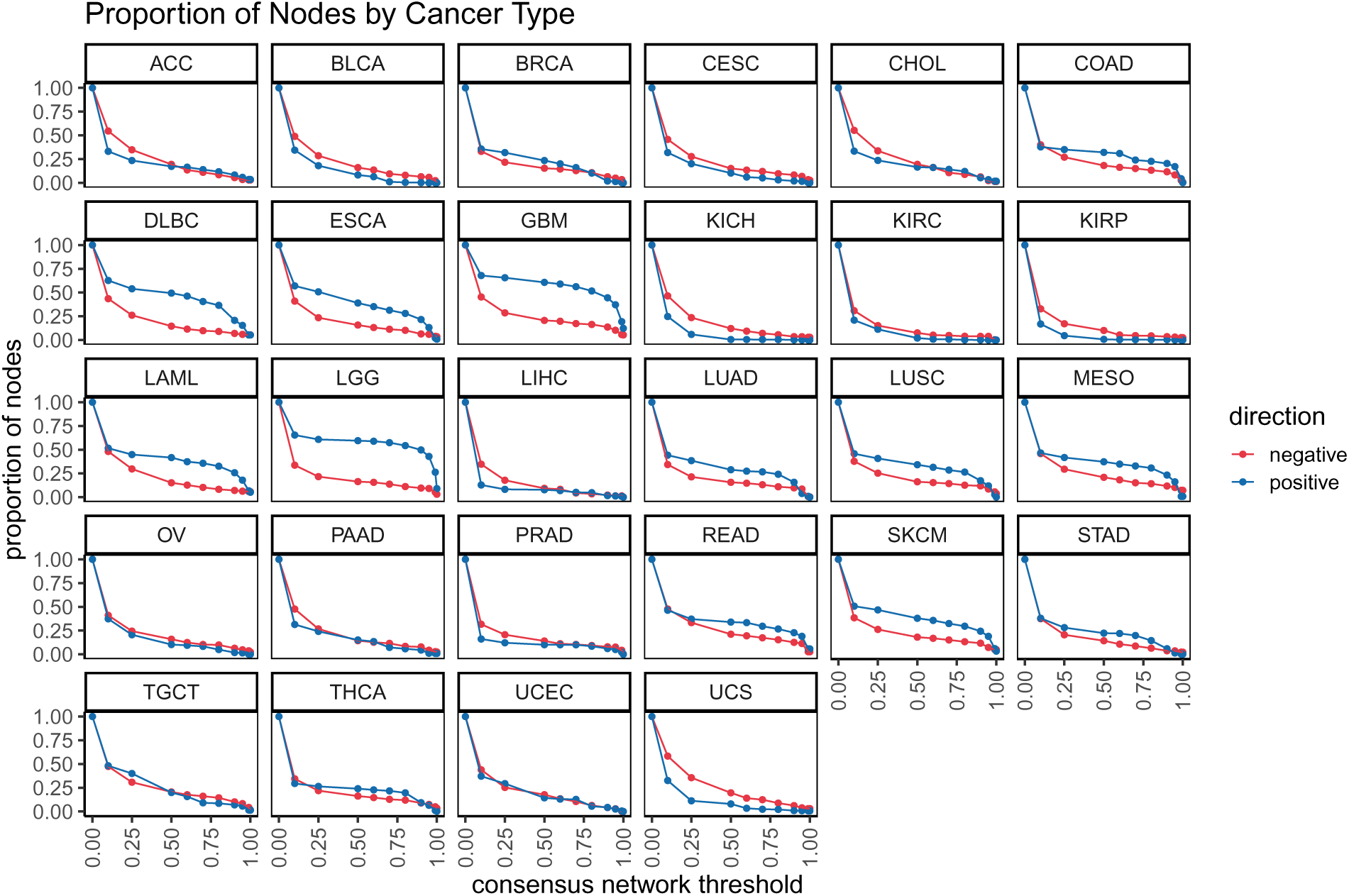
The line graphs show the number of nodes preserved for different consensus threshold for edge loss (negative, red) and edge gain (positive, blue) faceted by each cancer.

**Figure S3.**
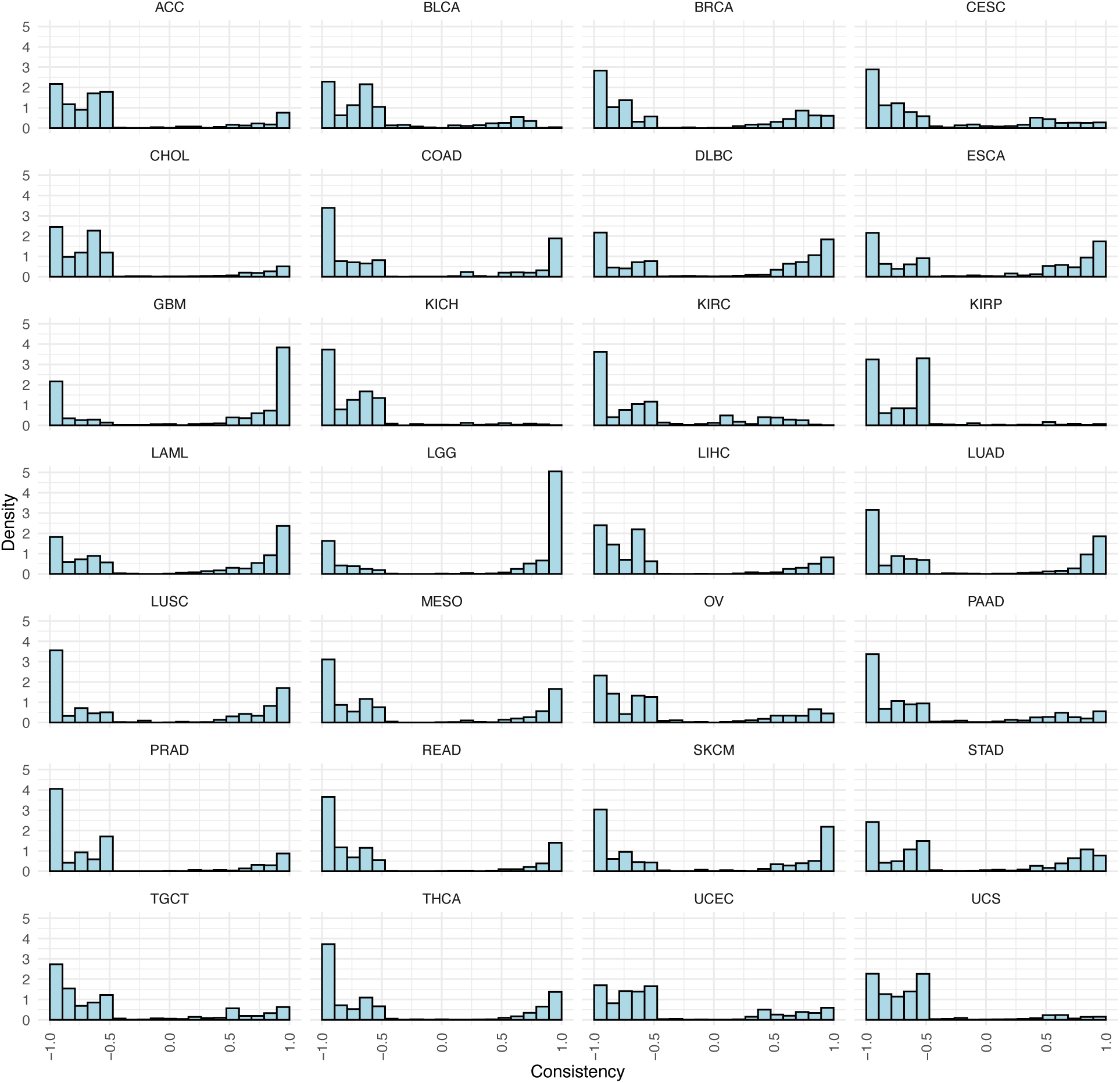
Density plots of edge consistency values faceted by cancer. Edges were filtered to only those that appear in 50% or more samples in that cancer.

**Figure S4.**
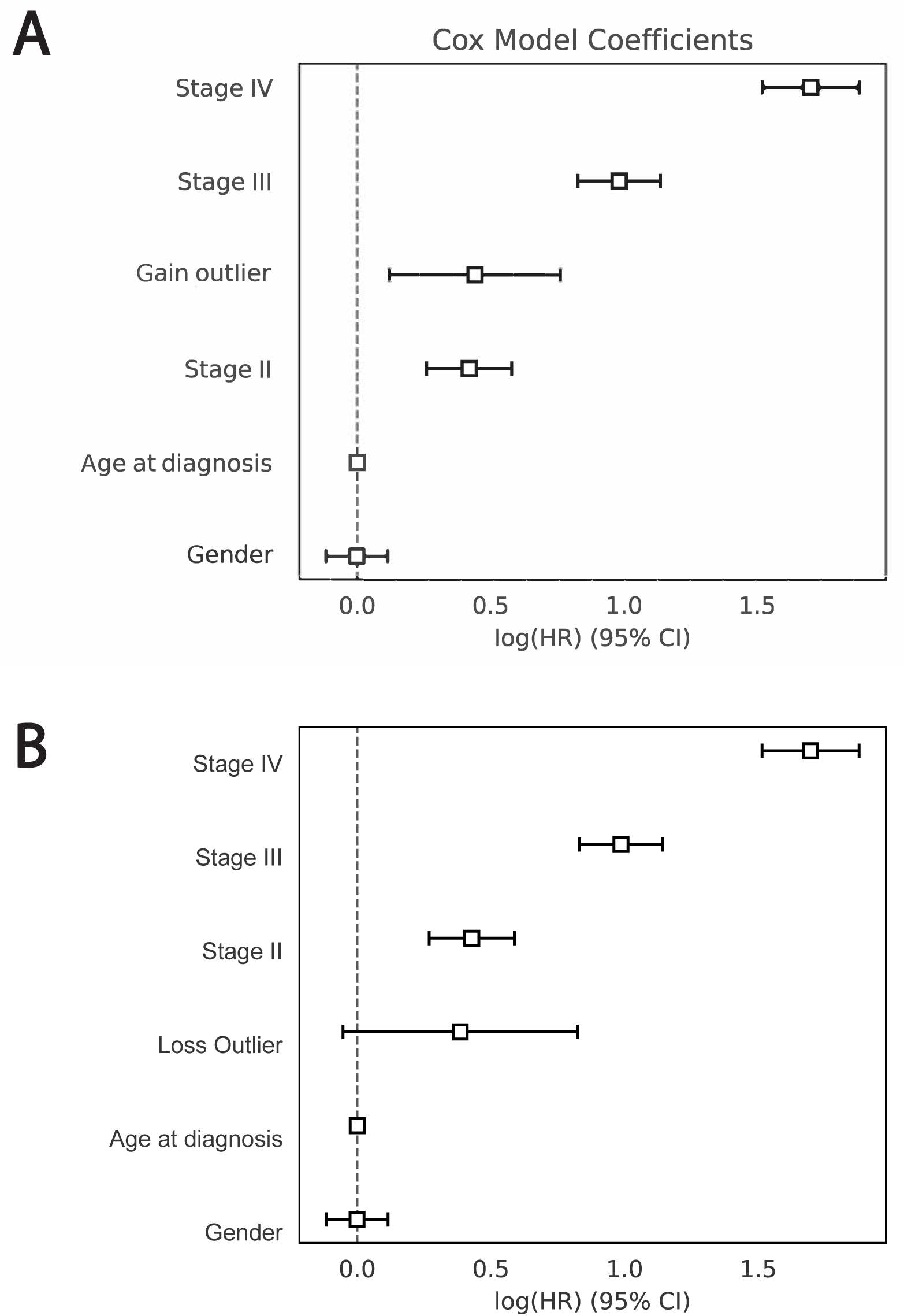
Forest plot of the multivariate Cox proportional hazards model coefficients, stratified by cancer type and adjusted for tumor stage, gender, age at diagnosis, and positive (A) and negative (B) outliers.

## Notes

https://zenodo.org/records/20767005

https://github.com/ylaboratory/splitpea-pancancer-analysis

### Competing Interest Statement

The authors have declared no competing interest.

### Funding

- Cancer Prevention and Research Institute of Texas: RR190065

